# CMAPLE 2: Fast and Accurate Phylogenetic Inference for Millions of Pathogen Genomes

**DOI:** 10.64898/2026.06.15.732229

**Authors:** Nhan Ly-Trong, Samuel Martin, Nick Goldman, Nicola De Maio, Bui Quang Minh

## Abstract

Phylogenetic analysis is essential to genomic epidemiology, for example in tracing the origin and evolution of SARS-CoV-2 variants during the COVID-19 pandemic. We previously introduced CMAPLE, a single-threaded implementation of the MAPLE algorithm designed for large-scale epidemiological genomic datasets. CMAPLE can reconstruct phylogenetic trees from up to one million SARS-CoV-2 genomes. Here, we present CMAPLE 2, a multi-threaded version of CMAPLE with parallel sample placement and subtree pruning and regrafting (SPR) search algorithms. CMAPLE 2 also reduces memory consumption by compressing data structures using multiple references along the tree instead of a single reference genome. It further implements two advanced models of highly site- and nucleotide-specific mutation patterns as observed in pandemic-scale genome data. Additionally, CMAPLE 2 parallelizes SPR-based Tree Assessment (SPRTA), an efficient and interpretable approach for assessing phylogenetic tree uncertainty, and supports ancestral state and mutation inference via mutation-annotated tree (MAT) reconstruction. When inferring a phylogeny from 500,000 SARS-CoV-2 genomes using 48 CPU cores, CMAPLE 2 reduces runtime from 5 days (with sequential CMAPLE) to 9 hours (a 13-fold speedup) while decreasing peak RAM usage from 11.1 GB to 7.3 GB. CMAPLE 2 can now reconstruct a tree of nearly four million SARS-CoV-2 genomes from scratch within 12 days using 41 GB of RAM, a task that the sequential CMAPLE and MAPLE cannot realistically complete. CMAPLE 2 is applicable to many pathogen genome datasets and enhances our preparedness for future pandemics.

---

Phylogenetics studies the origins and evolutionary relationships among organisms. It plays a vital role in reconstructing various clades in the Tree of Life (Misof et al. 2014; Parks et al. 2018; Dombrowski et al. 2020; Redmond and McLysaght 2021; Stiller et al. 2024; Williamson et al. 2025). It is also crucial for epidemiology, especially in viral phylodynamics (Volz et al. 2013), which uses pathogen genomic data to study how pathogens evolve and spread over time. During the COVID-19 pandemic, phylogenetics was crucial in tracing the origin and global spread of SARS-CoV-2, tracking transmission, identifying key mutations, and detecting the emergence of new variants (Coronaviridae Study Group of the International Committee on Taxonomy of Viruses et al. 2020; Gonzalez-Reiche et al. 2020; Rambaut et al. 2020; Hodcroft et al. 2021; Vöhringer et al. 2021; Su et al. 2024). At its core, phylogenetic inference takes as input a set of aligned DNA or protein sequences and reconstructs a phylogenetic tree representing the evolutionary history of those sequences.

We previously introduced CMAPLE (Ly-Trong et al. 2024), a C++ implementation of the MAPLE phylogenetic inference algorithm (De Maio et al. 2023) designed for pandemic-scale epidemiological genomic datasets. CMAPLE was optimized for time and memory efficiency and can reconstruct phylogenetic trees from up to one million SARS-CoV-2 genomes (Ly-Trong et al. 2024). However, CMAPLE is a sequential algorithm, i.e., using only one CPU core, and cannot deal with rapidly growing volumes of pathogen genome data expected during recent or future pandemics. Taking SARS-CoV-2 as an example, public repositories such as NCBI (ncbi.nlm.nih.gov/nuccore) now host over 9 million unrestricted open access complete and near-complete genomes, and more than 17 million genomes are available under a controlled-access model at GISAID (gisaid.org). This highlights the need for more efficient and scalable methods, especially to improve preparedness for future pandemics.

Here we introduce CMAPLE 2, a high-performance computing version of CMAPLE with substantial improvements in speed, scalability, memory efficiency, and functionality. Key advances include a parallel tree search algorithm utilizing multiple CPU cores, a compact data structure using sparse local references, two advanced site-specific variation models for evolutionary rates and nucleotide exchangeabilities, a parallel subtree pruning and regrafting tree assessment (SPRTA, De Maio et al. 2025b), and ancestral state and mutation inference via mutation-annotated tree (MAT, McBroome et al. 2021) reconstruction. CMAPLE 2 can now reconstruct a tree from nearly four million SARS-CoV-2 genomes (though its applications are not limited to this virus), exceeding the capabilities of both MAPLE and the sequential CMAPLE. These advancements are also available through the CMAPLE API library, facilitating integration into existing phylogenetic software. As an example, we integrated the parallel tree search and SPRTA algorithms into the widely used IQ-TREE 3 software (Wong et al. 2026).

In the following, we describe the key improvements and new features of CMAPLE 2, benchmark it against the sequential CMAPLE and the parallel version of MAPLE, and demonstrate its scalability on large datasets with millions of sequences. Throughout, we use CMAPLE to refer to the original sequential version and CMAPLE 2 to refer to the new parallel version presented in this paper. We also use “sequences” and “genomes” interchangeably, reflecting common usage in pathogen genomics.

## Parallel Tree Search Algorithm

The original CMAPLE (MAPLE) tree reconstruction algorithm comprises three stages: (1) building an initial rooted tree via iterative sample placements, (2) further optimizing the initial tree topology with subtree pruning and regrafting (SPR), and (3) re-optimizing all branch lengths. CMAPLE (MAPLE) uses a rooted tree structure. For non-reversible models, the root position is inferred by maximizing the tree likelihood, while for reversible models it can be arbitrarily assigned. The first two stages account for 97-98% of the total runtime.

Newer versions of MAPLE (De Maio et al. 2026) parallelized the SPR search stage 2 using multiple processes. While this reduces the run times of the second stage, the memory usage increases linearly with the number of processes, substantially limiting the scalability of MAPLE. To address these issues, we developed two parallel algorithms, presented below, for both stages 1 and 2 using the shared-memory multi-threading OpenMP library (Chapman et al. 2007). Since OpenMP is supported on most modern multi-core CPUs, users can benefit from parallelization on most, if not all, machines, from personal laptops and desktops to high-performance computing servers. Both algorithms can utilize up to *M* threads, where *M* is the number of available CPU cores.

### Parallel Sample Placement Algorithm

The sequential sample placement stage 1 in CMAPLE consists of two main operations: (1) searching for the optimal placement of each sample, and (2) inserting the sample into the tree. The former operation is independent across samples and therefore readily parallelizable, whereas the latter is more challenging to parallelize because each insertion will change the tree topology and affect subsequent placements. Our parallel algorithm exploits this structure by parallelizing the placement search while retaining a sequential insertion phase.

We first construct a ‘mini-tree’ sequentially using *S* samples (default *S* = 1,000). Construction of such small trees from SARS-COV-2 genomes with CMAPLE 2 usually takes only seconds and is not efficient to parallelize; however, for sequences with higher divergence a smaller value of *S* might be more efficient. The remaining samples are then divided into batches of size *M* × *K* (default *K* = 5), with placement searches and sample insertions performed one batch at a time. Batching is essential for maintaining placement accuracy: without it, all placements would be searched on the fixed mini-tree, which quickly becomes outdated as the tree grows. Processing samples in batches instead allows each batch to benefit from a progressively larger and more resolved tree constructed from previous batches. Each batch is handled in two phases. In the first phase, placements for all *M* × *K* samples are searched independently in parallel across *M* threads using dynamic scheduling. On average, each thread processes *K* samples. This is followed by a sequential refinement phase. Each sample’s placement is sequentially refined by starting a second narrower placement search from the *H*-step ancestor (the *H*-th ancestral node when traversing the tree upward to the root, or the root itself if it is reached before *H* steps; default *H*=5) of the initial placement node. This refinement phase is necessary because inserting a sample into the tree will modify it, thereby potentially affecting the exact optimal placement of subsequently inserted samples in the same batch. The sample is then inserted into the tree at the refined placement before continuing with the refinement and placement of the next sample in the current batch. The default values for the parameters *S, K*, and *H* were chosen to balance speed and accuracy on SARS-CoV-2 data, and these parameters can also be adjusted by users. We note that, as batch size (i.e., *M* ×*K* samples) depends on the number of threads *M*, the parallel sample placement algorithm may produce different trees when run with different numbers of threads.

### Parallel SPR Algorithm

The parallel SPR algorithm follows a two-phase strategy similar to the parallel sample placement. The first phase is the SPR search stage. Given the full initial tree relating all samples and constructed by the parallel sample placement algorithm with *M* threads, this phase seeks to improve the tree by performing SPR searches for all subtrees in parallel using dynamic scheduling. Unlike CMAPLE, which only considers regrafting a pruned subtree at the midpoint of a branch, CMAPLE 2 applies branch length optimizations that allow regrafting at any position along a branch to improve the accuracy. For each subtree, if at least one SPR move improving the tree likelihood is found, then we store the best SPR move (i.e., the one leading to the highest likelihood improvement) and its estimated likelihood improvement, but we do not apply yet any SPR move. After this parallel search completes, the stored candidate SPR moves are ranked by likelihood improvement.

In the second phase of the SPR algorithm, the stored candidate SPR moves are sequentially re-evaluated and applied to the tree starting from those with the highest likelihood improvement. Unlike sample insertions in the placement algorithm, which cause only local tree changes, SPR moves can significantly change the global tree topology, potentially invalidating likelihood improvements estimated for other candidate moves or moving their regrafting positions far from where they were originally found. Therefore, each candidate SPR move is re-evaluated by searching for the regrafting position from the root of the tree rather than from its *H*-step ancestor as in the sample placement algorithm. Usually the number of candidate (i.e. likelihood-improving) SPR moves found is very small compared to the total number of subtrees (because of the quality of the initial tree built by the parallel sample placement algorithm), making this sequential step relatively fast.

After each candidate SPR move is applied, all nodes whose partial likelihoods are changed are marked as “dirty”. Once the sequential phase completes, the parallel SPR algorithm is repeated with the SPR search restricted to subtrees rooted at dirty nodes. This process continues until the total likelihood improvement of the sequential phase becomes marginal (<1) or no dirty nodes remain, thereby completing one full iteration of the parallel SPR algorithm. By default, two such iterations are performed successively.

Several existing tools include parallel approaches for large-scale phylogenetic inference. UShER-sampled (Hinrichs et al. 2024) performs a similar parallel sample placement algorithm, but is based on maximum parsimony and has no batching like CMAPLE 2. For SPR-based optimization, matOptimize (Ye et al. 2022) also performs a parallel SPR search algorithm, but it uses the multiprocessing MPI library (Gropp et al. 1998). RAxML-NG (Kozlov et al. 2019) and IQ-TREE parallelizes likelihood calculations across alignment sites, which scales well for long phylogenomic alignments. For pandemic-scale datasets where taxa significantly outnumber sites, parallelizing across samples or subtrees, as in UShER-sampled, matOptimize, and CMAPLE 2, is more effective.

### Compact Data Representation using Multiple Local Reference Genomes

Taking advantage of the high sequence similarity of typical pathogen data, earlier versions of CMAPLE and MAPLE represent each genome as a set of differences relative to a (single, global) reference sequence (De Maio et al. 2023; Ly-Trong et al. 2024). This representation can reduce input size by more than 100-fold, enabling efficient phylogenetic reconstruction on large-scale pathogen data (De Maio et al. 2023).

Over time, as pathogens continue to evolve, within-outbreak divergence from the initial genome increases and progressively undermines the performance of reference-based representation. To address this limitation, CMAPLE 2 implements a more compact data structure based on a hierarchical system with multiple local references distributed across the tree (De Maio et al. 2026). These local references are generated automatically and require no additional input from the user. Each local reference is applicable within a subtree, i.e., genomes are encoded as differences with respect to their closest ancestral local reference, reducing the number of differences that must be recorded and enabling more efficient storage, likelihood evaluation, and tree rearrangement operations.

Maintaining local references incurs overhead from tracking the appropriate reference for each genome considered during likelihood calculations and updating local references during tree rearrangements. We found that best results were achieved by limiting overhead and using only a relatively sparse sampling of nodes selected as local references (De Maio et al. 2026). A node is assigned a local reference if it has at least 50 descendant branches with non-zero lengths and accumulates at least two mutations relative to its most recent ancestral reference (i.e., the nearest reference on the path from that node to the root). During sample placement, these criteria determine when new local references are created. During SPR optimization, local references are moved together with their pruned subtrees, and their stored substitutions are updated following regrafting. However, the reference-assignment criteria are not re-evaluated during this stage, and no new local references are introduced. These criteria were optimized for SARS-CoV-2 data and can be adjusted by users if useful.

### Rate Variation Models

Evolutionary rates vary across genomic positions due to differences in functional constraints, selective pressures, and mutation processes (Kimura and Ohta 1974; Yang 1996). A standard approach to model this heterogeneity is the Gamma model (Yang 1994a), which assumes that site rates follow a Gamma distribution (typically discretized into a small number of categories). However, it is hard for this approach to scale to large datasets and to account for the highly site- and nucleotide-specific mutational patterns observed in SARS-CoV-2 (De Maio et al. 2021, 2025a, 2026).

To address this limitation, we developed two complementary models. The first is a site-specific rate model that assigns an independent scaling factor of substitution rates to each site (De Maio et al. 2026). The second is a Site-Specific Matrix (SSM, Martin et al. 2026) model that assigns an independent UNREST substitution matrix (Yang 1994b) to each site. This enables both site- and nucleotide-specific variation: for example, it allows a substitution such as G→T to be highly recurrent at a given site, while G→C at the same site is not. These models are highly parameter-rich, and thus primarily suited to datasets with many genomes.

### SPR-based Tree Assessment (SPRTA)

Assessing the uncertainty of inferred phylogenies is essential for interpreting evolutionary and epidemiological analyses (Lemoine et al. 2018). CMAPLE 2 implements and parallelizes the recently proposed SPR-based Tree Assessment (SPRTA) method (De Maio et al. 2025b). Unlike classical branch support measures such as Felsenstein’s bootstrap (Felsenstein 1985), which evaluate confidence in clade existence, SPRTA assesses the probability that a sample or lineage evolved from a particular ancestor rather than from other alternative ancestors in the tree. While Felsenstein’s bootstrap is computationally prohibitive at pandemic scale, SPRTA remains feasible for such datasets. For each branch, SPRTA computes an approximate posterior probability for multiple plausible evolutionary origins by comparing the likelihood of the inferred tree against alternative SPR topologies. Because these SPR moves are already evaluated during CMAPLE’s SPR search, SPRTA incurs negligible additional computational cost and gains full benefit from CMAPLE 2’s parallelization. Applying an SPR move affects the likelihoods of nearby nodes, making their SPRTA scores outdated. These scores are automatically recomputed during the subsequent SPR search on dirty nodes (see Parallel SPR Algorithm), so that the final SPRTA values are updated to better reflect the new tree topology.

### Mutation Annotated Trees (MATs)

CMAPLE 2 can estimate mutations occurring along the branches of the inferred tree. For each branch, it computes the posterior probability of possible mutations that might have occurred on the branch, conditional on the inferred substitution model and tree, and the genetic data. The resulting mutation-annotated tree (MAT: Turakhia et al. 2021) is output in NEXUS format, with inferred mutations and their posterior probabilities annotated on each branch.

### Benchmark of CMAPLE 2 against CMAPLE and MAPLE

We benchmarked CMAPLE 2 (v2.0.0) against CMAPLE (v1.0.1) and the latest parallel version of MAPLE (v0.7.5) on a server equipped with two Intel(R) Xeon(R) Gold 6252 CPUs (2.10 GHz, 2 × 24 cores). CMAPLE was evaluated using a single CPU core, CMAPLE 2 using 16 and 48 cores, and MAPLE using 16 cores only (due to memory constraints). MAPLE was executed with pypy3.9-v7.3.9 (https://pypy.org/) to maximize performance. For testing data, we sampled between 100K and 3.8M real SARS-CoV-2 genomes assembled with Viridian (Hunt et al. 2026). All analyses employed the UNREST substitution model (Yang 1994b).

CMAPLE 2 is substantially faster and consumes much less memory than CMAPLE and MAPLE (Figure 1). To reconstruct a tree from 500K SARS-CoV-2 sequences, CMAPLE using a single CPU core required 116.7 hours and 11.1 GB RAM, whereas CMAPLE 2, using 48 CPU cores, required only 9.1 hours (13x faster) and 7.3 GB RAM (34% lower memory usage). Compared with MAPLE using 16 CPU cores, which required 40.6 hours and 343.1 GB RAM, CMAPLE 2 was 4.5x faster while reducing memory usage by 98%.

**Figure 1.**
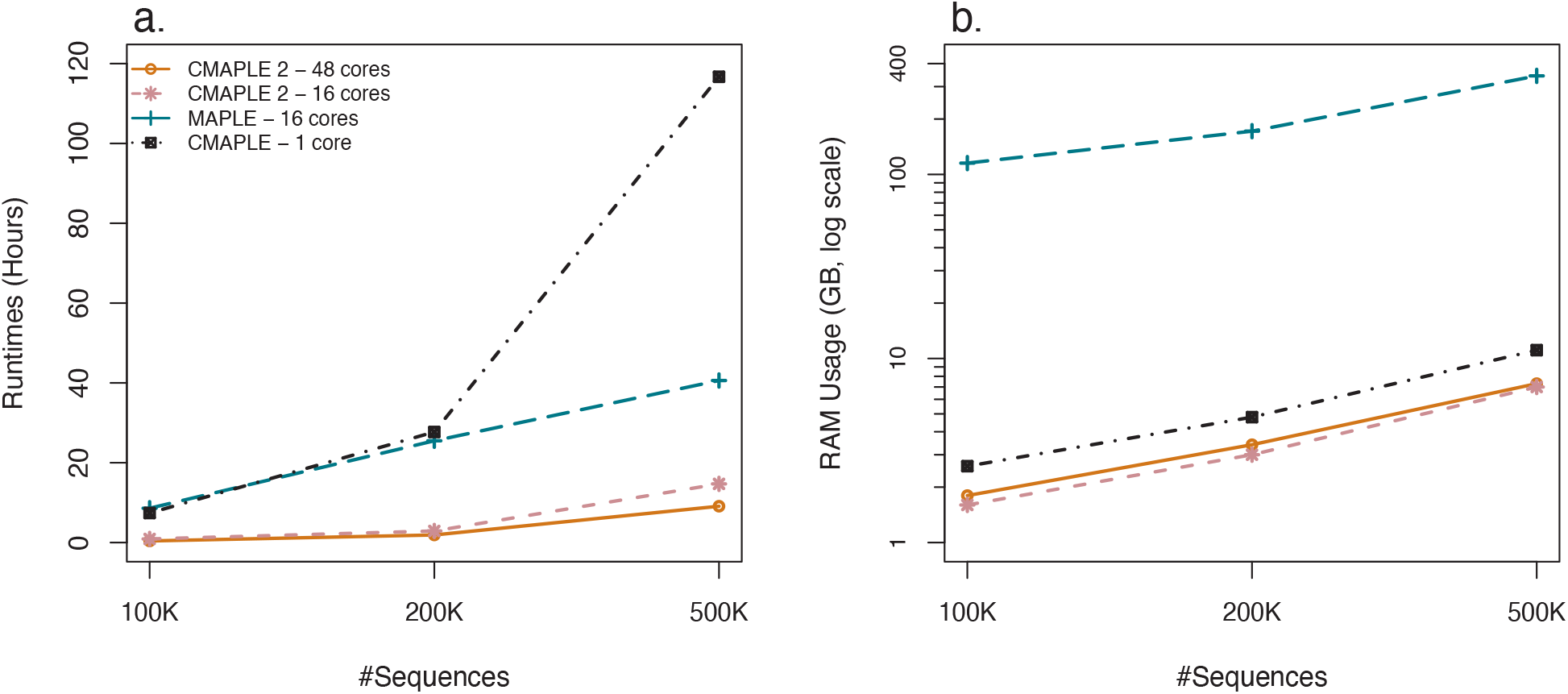
Runtimes (a) and log-scale peak memory consumption (b) of CMAPLE, CMAPLE 2, and MAPLE when analyzing 100K, 200K, and 500K randomly sampled SARS-CoV-2 genomes.

### Performance of CMAPLE 2 on Millions of Pathogen Genomes

By adopting a shared-memory multi-threading paradigm, CMAPLE 2 avoids the excessive memory overhead of MAPLE, enabling CMAPLE 2 to handle much larger alignments. We further evaluated the scalability of CMAPLE 2 on datasets comprising 1M, 2M, 3M, and 3.8M real SARS-CoV-2 genomes. Runtime and memory usage of CMAPLE 2 using 48 CPU cores are reported in Table 1. CMAPLE 2 reconstructed a phylogeny from nearly 4M genomes from scratch in 12 days using 41 GB of memory, a task that neither the sequential version of CMAPLE nor the parallel version of MAPLE could complete.

**Table 1.** Runtime and peak memory consumption of CMAPLE 2 using 48 CPU cores for phylogenetic reconstruction from 1M, 2M, 3M, and 3.8M real SARS-CoV-2 genomes.

| Data size | Reference genomes | Runtime (days) | Peak memory consumption (GB) |
| --- | --- | --- | --- |
| 1M | global | 2.34 | 28.59 |
| 1M | local | 1.81 | 13.73 |
| 2M | local | 4.13 | 23.26 |
| 3M | local | 11.73 | 34.38 |
| 3.8M | local | 12.32 | 41.00 |

We also assessed the advantage of using multiple local references over a single global reference within CMAPLE 2. When reconstructing a tree from one million SARS-CoV-2 genomes using 48 CPU cores, multiple local references reduced runtime from 2.34 to 1.81 days (1.29x speedup) and memory consumption from 28.59 GB to 13.73 GB (52% reduction).

### Software Validation

We validated our implementation by comparing the log-likelihoods of trees inferred by CMAPLE (v1.0.1), CMAPLE 2 (v2.0.0), and MAPLE (v0.7.5) using five independent random subsamples of 100K SARS-CoV-2 genomes. The aim was to verify that the speedup gained in CMAPLE 2 does not sacrifice accuracy, i.e., CMAPLE 2 reconstructs trees with likelihoods not lower than those of CMAPLE and MAPLE. Additionally, we evaluated CMAPLE 2 using both global and local reference sequences, with the expectation that the two settings would produce trees with comparable likelihoods. For comparisons, log-likelihoods were recomputed using IQ-TREE (v3.0.1).

Across all alignments, CMAPLE 2 often produced trees with higher likelihoods than MAPLE and consistently outperformed CMAPLE (Suppl. Table 1). This is likely because CMAPLE 2 performs more thorough SPR optimisation, with finer-grained branch length optimisation during the SPR search. Furthermore, trees inferred using global and local references yielded comparable likelihoods (Suppl. Table 1), with minor differences likely due to approximations introduced in the local reference implementation to reduce computational complexity.

The CMAPLE 2 rate variation models are validated in Martin et al. (2026). Additionally, we compared SPRTA supports produced by CMAPLE 2 and MAPLE on a 100K-genome SARS-CoV-2 phylogeny inferred as described above. The two sets of support scores have a Pearson correlation coefficient of 1.0, and a root mean square difference of 0.0013 (Suppl. Figure 1). To validate the MAT reconstruction, we compared CMAPLE 2 and MAPLE on an alignment of 100K SARS-CoV-2 genomes. For each mutation type (i.e. mutation identity and location; e.g., G21307C, a G→C mutation at site 21307), we evaluated (i) the expected number of mutations, defined as the sum of the posterior probabilities for the mutation type across the entire tree, and (ii) the number of branches with non-negligible (>1%) posterior probability for the mutation type. Both metrics showed strong concordance between CMAPLE 2 and MAPLE, with Pearson correlation coefficients of 1.0 and root mean square difference of 0.119 and 0.222 respectively (Suppl. Figure 2).

We assessed the code quality of CMAPLE 2 using SoftWipe (Zapletal et al. 2021), achieving an overall program absolute score of 7.1/10, ranking seventh among 49 computational tools (Zapletal et al. 2021). All datasets and testing scripts used in this project are provided in the Supplementary Material.

## Conclusion

CMAPLE 2 introduces substantial improvements over its earlier version, adding a scalable parallel tree search algorithm, an enhanced data structure, advanced evolutionary models, and new features for assessing phylogenetic uncertainty and inferring mutation histories. By employing local references, CMAPLE 2 should scale to more diverged datasets than the original CMAPLE. Together, these developments enable efficient pandemic-scale phylogenetic inference and support more accurate genomic epidemiological analyses. The parallel tree search and SPRTA assessment have also been integrated into IQ-TREE 3 (Wong et al. 2026) via the CMAPLE APIs, which are designed to facilitate rapid adoption by the research community.

Further improvements could be achieved by fine-tuning the default parameters, which were originally optimized on SARS-CoV-2 data. Dynamically and automatically adapting these parameters based on the input data at hand, i.e., when it is not SARS-CoV-2, would be desirable, and could further improve both computational efficiency and accuracy across a broader range of pathogens.

Several directions exist to potentially further improve the scalability of CMAPLE 2. First, parallelization could be extended to distributed computing environments using the Message Passing Interface (MPI, Gropp et al. 1998). This would enable computations to be distributed across multiple nodes in high-performance computing clusters, allowing phylogenetic inference at scales beyond the capacity of a single machine. Second, GPU acceleration is also a promising avenue. Reconstructing a tree from 3.8M SARS-CoV-2 genomes required 41 GB of RAM, which fits the memory capacity of modern high-end GPUs. This opens the possibility of leveraging GPUs to accelerate tree search and further enhance the scalability of CMAPLE 2.

## Acknowledgments

We thank Robert Lanfear for his valuable comments and discussions. We used ChatGPT to assist in spelling checks and grammar corrections during the preparation of this manuscript.

## Data Availability

CMAPLE 2 (v2.0.0) is released under the GPL-2.0 open-source license with precompiled binaries and source code at https://github.com/iqtree/cmaple. The user manual and command reference are available at https://github.com/iqtree/cmaple/wiki/User-Manual. CMAPLE API documentation is available at http://iqtree.github.io/doc/cmaple. All the scripts necessary to reproduce the analyses reported in this study can be accessed through Zenodo at https://doi.org/10.5281/zenodo.20620422.

---

**Supplementary Table 1.** Log-likelihoods (computed by IQ-TREE) of trees inferred by CMAPLE, CMAPLE 2 (with global and local references), and MAPLE from five alignments of 100K real SARS-CoV-2 genomes subsampled from the Viridian alignment (Hunt et al. 2026). The highest likelihood for each alignment is shown in **bold**. (*) indicates whether global or local references yield the higher-likelihood tree.

|  | CMAPLE | CMAPLE 2 |  | MAPLE |
| --- | --- | --- | --- | --- |
|  |  | Local references | Global reference |  |
| 100K_1 | -2,354,689 | -2,354,677 | **-2,354,499\*** | -2,354,641 |
| 100K_2 | -2,351,253 | **-2,351,206\*** | -2,351,245 | -2,351,346 |
| 100K_3 | -2,356,216 | -2,356,192 | **-2,356,149\*** | -2,356,246 |
| 100K_4 | -2,352,490 | -2,352,382* | -2,352,480 | **-2,352,341** |
| 100K_5 | -2,349,511 | -2,349,359 | **-2,349,327\*** | -2,349,385 |

**Supplementary Figure 1.**
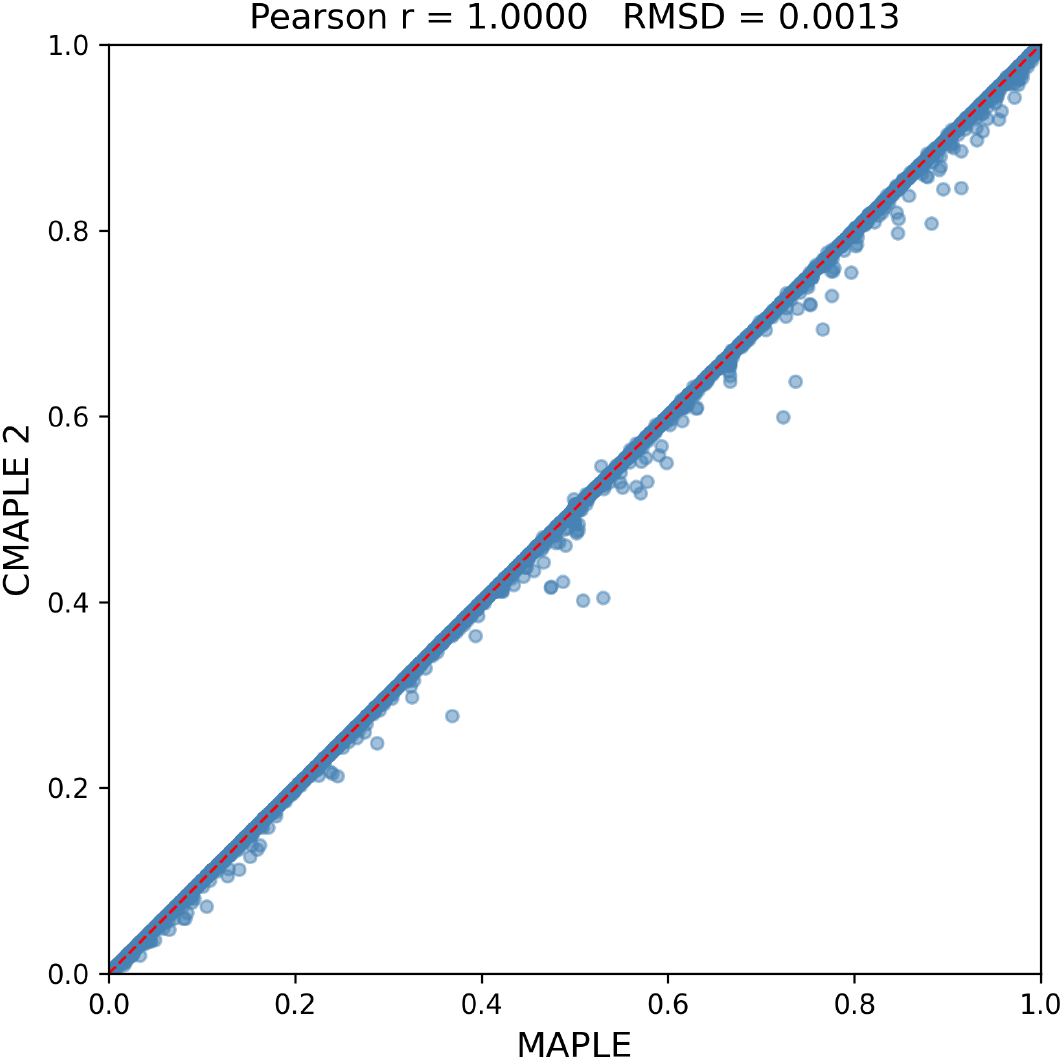
Scatter plot comparing SPRTA support scores computed by CMAPLE 2 and MAPLE on a 100K-genome SARS-CoV-2 phylogeny. Each point indicates the scores assigned to one branch of the phylogeny by the two methods. The two methods showed a strong concordance, with a Pearson correlation coefficient of 1.0.

**Supplementary Figure 2.**
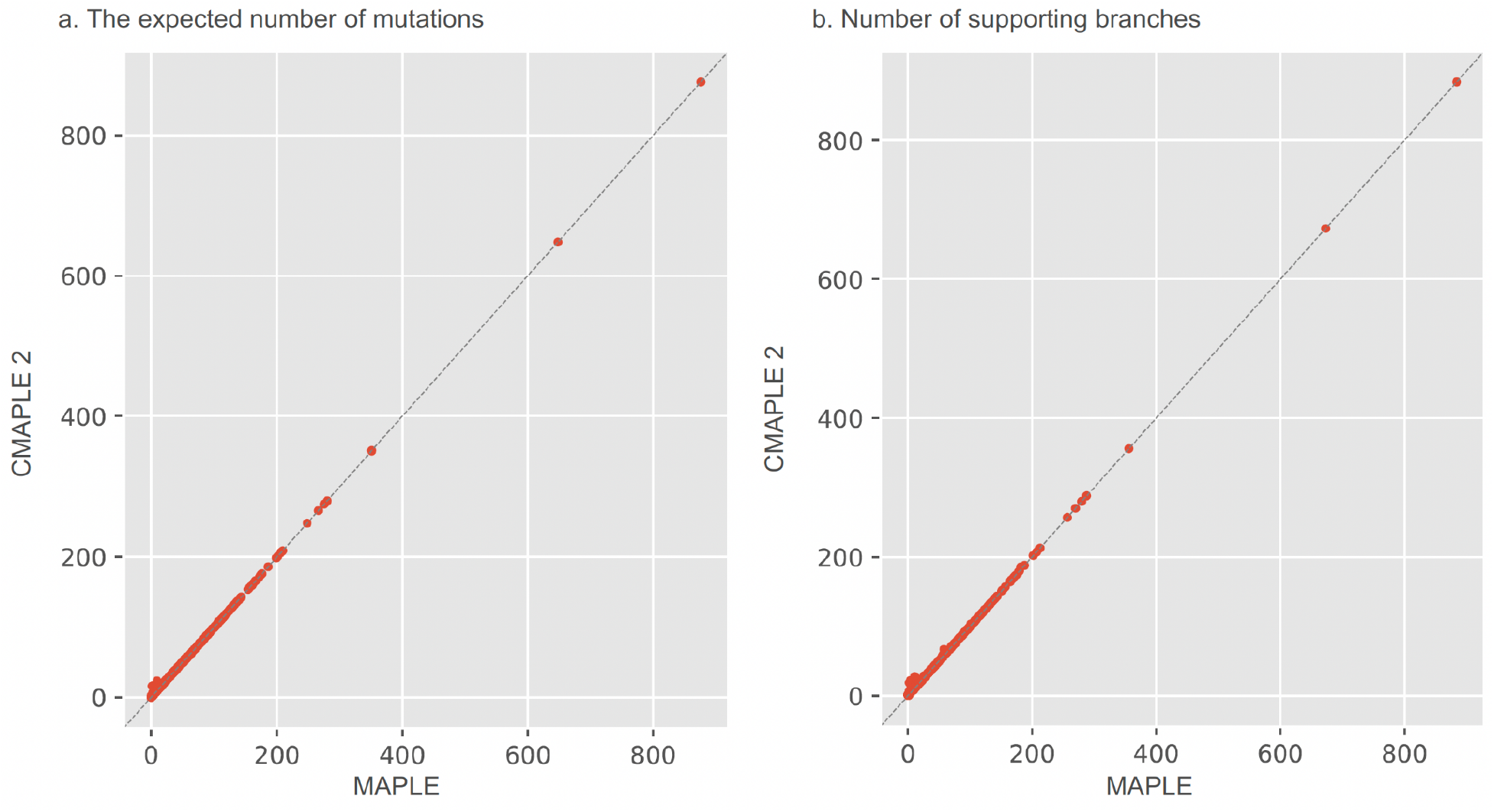
Comparison of MATs inferred by CMAPLE 2 and MAPLE on an alignment of 100K SARS-CoV-2 genomes. For each plausible mutation at a branch, both tools compute a posterior probability. (a) Expected number of mutations: the sum of the posterior probabilities for a given mutation type across all branches. (b) Number of supporting branches: the number of branches that support a particular mutation type (i.e. branches with a posterior probability of >1% for that mutation).

